# A method to study honeybee foraging regulatory molecules at different times during foraging

**DOI:** 10.1101/2021.05.28.446211

**Authors:** Asem Surindro Singh, Machathoibi Chanu Takhellambam

## Abstract

The foraging of honey bees is one of the most well organized and admirable behaviors that exist among social insects. In behavioral studies, these beautiful insects have been extensively used for understanding time-space learning, landmark use and concept of learning etc. Highly organized behaviors such as social interaction and communication are systematically well organized behavioral components of honeybee foraging. Over the last two decades, understanding the regulatory mechanisms underlying honey bee foraging at the cellular and molecular levels has been increasingly interested to several researchers. Upon the search of regulatory genes of brain and behavior, immediate early (IE) genes are considered as tool to begin the investigation. Our two recent studies, have demonstrated three IE genes namely Egr-1, Hr38 and kakusei having a role in the daily foraging of bees and their association with learning and memory during the foraging. These studies further evidence that IE genes can be used as a tool in finding the specific molecular/cellular players of foraging in honey bees and its behavioral components such as learning, memory, social interaction, social communication etc. In this article we provide the details of the method of sample collection at different times during foraging to investigate the foraging regulatory molecules.

## Introduction

Honey bee foraging consists of several behavioral components that include search of food, identification and memorization of food source location, carrying and storing of food, interaction and communication etc (Frisch 1965; Seeley 1995; Singh 2019; Singh et al., 2020; Singh et al., 2018;). Inside the artificial bee house, honey bees flew several times to and fro carrying pollen/nectar from the feeder to the hive. All of these behaviors can be observed by feeding the bees few meters away from the hive inside the bee house. Generally, cap honey bees or European honey bee *Apis mellifera* is widely used for understanding behavioral dynamics of social insects. Though it looks small and tiny, this insect displays incredible means to communicate each other. During foraging honey bees of the same colony share information and communicate each other through waggle dance which a typical movement of the honey bees that looks like numerical figure eight. For the first time this waggle dance of bees was translated by Austrian ethologist Karl Ritter von Frisch and he received Nobel Prize in Physiology or Medicine in 1973 for his incredible effort towards investigating the sensory perceptions in honey bees (Michelsen 2003; Von Frisch 1993; Riley et al 2005; Seeley et al 2006).

So far, the foraging behavior of honey bees has been extensively studied and available data are plenty. This has opened an incredible research opportunities for understanding the regulatory mechanisms at various ways in honey bees. But the reports on the underlying mechanisms in the brain are very limited. However there is increasing interest on this direction in the recent years. Understanding cellular and molecular regulators of honey bee foraging, would not confined only within insects, but also can offer some guidelines/clues towards unwinding the complexity of neural circuitry and molecular underpinnings in the higher animals and humans (Singh 2014). Because humans and other large animals have many similar behavioral features with honeybees, and there is sequence homology in the genes across various species.

Upon the search of foraging regulatory genes in the brain, it is a promising way to begin with IE genes. Because IE genes are well known neural markers. The IE genes had been found to have persisting roles from the first stages of brain development unto the adulthood showing possible inherent features in the everyday brain activity (Loebrich and Nedivi, 2009). It also showed with dramatic roles in phenotypic changes that occurred in neurons (Dijkmans et al., 2009). There are many IE gene encoded transcription factors that are rapidly induced within the neurons and some were delayed (Friedman et al., 1992; Kaczmarek 1993) in response to different stimuli and cellular environments that result to neuronal capacities with short- and long-lasting phenotypic changes (Loebrich and Nedivi, 2009; Hughes and Dragunow, 1995). Following the stimulation, early response neurons reacts from milliseconds to minutes and this involves 1st and 2nd messenger systems and phosphatases, whereas late response persists from hours to days and even to permanent changes coupled with gene expression changes (Hughes and Dragunow, 1995; Clayton 2013). The late response neurons were found to link with learning, memory, sensitization processes and drug tolerance habits etc. (Hughes and Dragunow, 1995; Clayton 2013). Remarkably, during the process of nerve stimulation, the IE genes have been noted as first activated genes that link to membrane events and nucleus (Beckmann and Wilce, 1997). Therefore, regulation in the IE gene expression level is considered to be the first part in general neuronal response to a natural stimulus. And thus, it is a promising way to start with IE genes in the planning of experimental strategies from scratch towards finding specific molecular and neuronal pathway leading to a specific behavior.

In our two recent studies, we have found three IE genes, *Egr-1, Hr-38* and *kakusei* involvement in the daily foraging of honey bees as well they possible role in learning, memory processing during foraging [Singh et 2018; Singh et al 2020]. Moreover, possible role of downstream genes of *Egr-1*, ecdysone receptor (*EcR*), dopamine/ecdysteroid receptor (*DopEcR*), dopamine decarboxylase (*Ddc*) and dopamine receptor 2 (*Dr2*) were also revealed. Subsequently, involvement of *Hr38, EcR* and *DopEcR* which are part of ecdysteroid signalling pathway, in learning and memory processes, have been also indicated (Singh et al 2018). We have performed the sample collections/behavioral experiments while the bees were on the process of continue foraging and such an experiment was not before. In this article we describe the method of sample collection and the molecular experiments in detail.

## Materials and Methods

### Behavioral experiment and sample collection

European honey bee species (also known as cap honey bees) *Apis Melifera* colonies were purchased from the local bee keepers in Bangalore, Karnataka, India. The bee colonies were placed inside the bee house of the institute, National Centre for Biological Sciences (NCBS), Tata Institute of Fundamental Research (TIFR), Bangalore, India. *Apis Melifera* is one of the most common and widely domesticated honey bee species in the world. Therefore this honey bee species is not endangered or threatened and available almost every place in the world. Behavioral test was performed inside the bee house of NCBS which is an outdoor flight cage. The bee house is 12m in length, 5m in height and 2.5m in width. The bees were fed with pollen in a green plastic place and 1 M sucrose solution in a yellow plastic plate. The distance of feeders from the beehive was 10m and the two feeders were kept 1.5m apart from each other. The bees were fed everyday from 14.00 to 17.00 hours.

Sample collection was started after the foraging bees had learned and adapted about the location of feeders and had visited the feeders for several days. The collected samples were subjected to gene expression profiling and for this only nectar foragers were collected. The procedures were briefly described in our two previous articles (Singh et al 2018; Singh et al 2020) and in this article we described the method in detail.

### Sample Collection procedure and grouping

#### Sample collection during foraging

For the 0min group sample, the first arriving foragers at the feeder plate were caught before presenting the sucrose solution on plate and the caught bees were immediately flash frozen in liquid nitrogen. For catching the bees, 50 mL falcon tubes with tiny pores were used. As soon as the first collection over, sucrose solution was poured and some first arriving foragers were gently marked using Uni POSCA Paint Markers (Uni Mitsubishi Pencil, UK) on the head while they are drinking sucrose solution and time count was immediately started. The marked bees that arrived on the feeder during their repeated trips and they were gently caught at a series of different time points with 15 min intervals up to 2 hours. The time points were 15min, 30min, 45min, 60min, 75min, 90min, 105min and 120min. After catching, the bees were immediately flash frozen in liquid nitrogen. About 24 bees were collected in a day from 14.00 hours to 16.00 hours, i.e., 1–2 bees for each time point and continued in following days until 5 bees were obtained at each time point group. The collected samples were stored at - 80^0^C for further experiment.

#### Sample collection for before and after foraging groups

For collecting the before-foraging samples, some first arriving bees on the feeder at 14:00 hour were paint-marked (in the same way as above) and collected in the following day morning at 09.00 hours inside the hive, before they started flying out from the hive for foraging. In the case of after-foraging samples, the bees which were paint-marked were caught in the hive in the evening at 18.00 hours of the same day of paint marking after the bees finished foraging. Gentle care had been taken always during the collection procedure in order not disturb the bees normal behavior and to avoid inducing stress phenomena; in this way minimal interactions between the collector and the bees could be accouned (Abbott and Dukas 2009; Bateson et al 2011; Nieh 2010). The caught bees were immediately flash frozen in liquid nitrogen and stored at -80^0^C for further processing. In this case 5 bees in each group were collected at the same time on the same collection day.

#### Sample collection for food un-reward group

This group consisted only the foraging bees collected on the empty feeder plate and the collection were done for 1 hour at four time points with the intervals of 15min. The 0min samples were collected at 14:00 hours on the empty feeder plate and immediately followed by paint-marking of some bees (in the same way as above) for the collection of the four other time points. Since bees had to be collected without food reward at all four time points, a simple trick was needed to be applied. A small amount of 1 M sucrose solution was presented on the plate to let the bees continue foraging on the feeder but a little portion was allowed to be accessible to the bees for drinking by covering sucrose solution with a transparent bowl. This way the marked bees were stopped from drinking sucrose solution and made sure that they did not touched the sucrose solution. And while some unmarked bees were drinking, the paint-marked bees were caught as soon as they landed on the feeder plate. About 1-2 bees were collected in each day and the collection was done from 14.00 to 16:00 hours and each group had 5 bees at least. The bees were immediately flash frozen in liquid nitrogen as soon as they were caught and stored at -80^0^C for further experiments.

### Gene expression profiling

#### Brain dissection

The frozen bees at -80°C were removed and lyophilized at -50°C with vacuum at 0.420 mBar for 20min, using lyophilizer (Freeze Zone1 PlusTM 4.5 liter cascade Freeze Dry System, Labconco Corporation, Kanas City). The bee head was placed in a glass chamber containing 100% ethanol placed on a dry ice platform and the brain dissection was performed under a light microscope. Soon after the dissection, the whole brain was immediately placed into a 1.5 mL eppendorf tube placed on dry ice, and 500 μL Trizol (Trizol Reagent, ambion RNA, life technology) was added. Thus prepared sample was ready for total RNA preparation. The same procedure was followed for every bee brain dissection.

#### RNA preparation and cDNA conversion

The frozen sample was thawed placing on ice and the brain was homogenized using an electronic homogenizer (Micro-Grinder Pestle Mixer, RPI Research Products International) with pestle (Micro-Tube Sample Pestles, Research Products International). By centrifugation at 10000g for 5min at 4^0^C, the total RNA, protein, DNA and cell debris fractions were separated. The upper clear fraction containing RNA was removed gently, leaving the DNA, tissue debris and the protein fractions. And thus the total RNA was extracted. Then cDNA was prepared from the total RNA, by using the kits supplied by Invitrogen (Thermo Fisher Scientific). Manufacturer’s protocol was followed in cDNA preparation.

#### Quantitative real time (qPCR)

The prepared cDNA in the above, from each brain was subjected to qPCR using a 7900HT Fast Real Time PCR System (Applied Biosystem, Singapore). The qPCR reaction mixture for each sample was prepared in 200μL microcentrifuge tubes with 10μl reaction volume that contained cDNA, oligonucleotide primers (Sigma Aldrich) specific to target genes and SYBR Green (KAPA Syber1 FAST PCR Master Mix (2X) ABI Prism1). The qPCR cycles followed Applied Biosystem protocol. Rp49 gene was used as endogenous control in every qPCR run. The details of the target genes and the oligonucleotide primers are provided in Table 1.

**Table 1.**
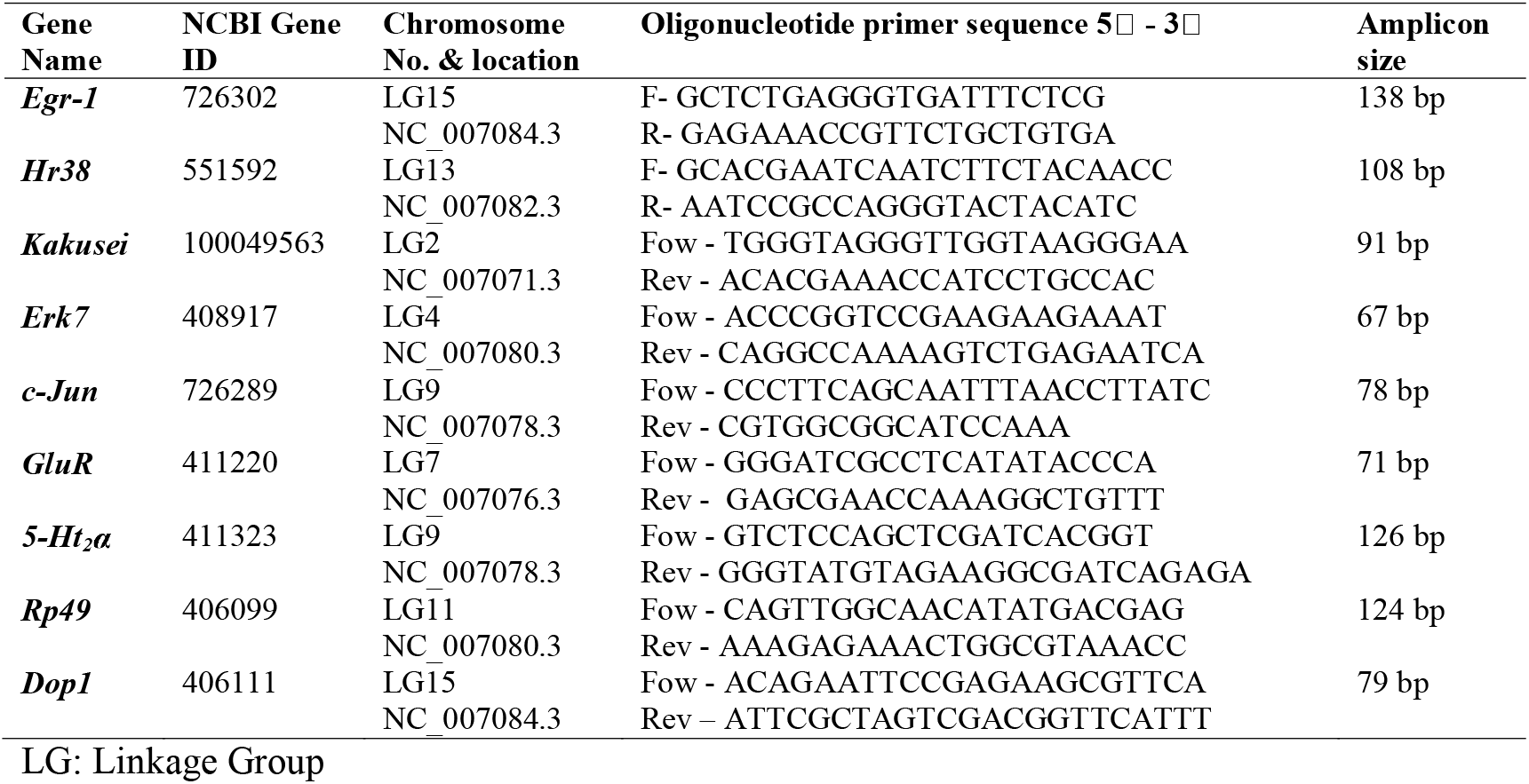
Gene locations, oligonucleotide primers and the qPCR product amplicon sizes.

### Statistical analysis

The relative gene expression level was calculated using the relative standard curve method with the help of SDS 2.4 software provided with the 7900HT Fast Real system. The standard deviation was calculated following Applied Biosystem’s ‘Guide to performing relative quantification of gene expression using real-time quantitative PCR’. The fold changes at all the times points was determined relative to time t0, and the statistical significance was examined using one-way ANOVA with Turkey-Kramer post-hoc multiple comparison test and the analysis was carried out with the help of GraphPad InStat software (Motulsky 1999). Normal distribution of each comparing group was tested using the D’Agostino & Pearson omnibus normality test.

## Results

### IE genes, *Egr-1, Hr-38* and *Kakusei* expression during foraging, before foraging and after foraging

In this study, we have combined most of our recent published data in two different journals (Singh et al., 2018; Singh et al., 2020). A total of nine genes have been considered including a house keeping gene *Rp49*. The eight genes are *Egr-1, Hr38, Kakusei, c-Jun (Jra), Erk7, GluR, 5-Ht2*α, *DopR* and further details of these genes are provided in Table 1. Among the four IE genes *Egr-1, Hr38, Kakusei, c-Jun (Jra)*, we observed that *Egr-1, Hr38 and Kakusei* were found to have significant transient over expression during the reward foraging in two hours. The results are summarized in Figure 1. The other four genes *Erk7, GluR, 5-Ht2*α, *DopR* as well as *c-Jun (Jra)* were found to have no difference in their expression during foraging, shown in Figure 2. Further, there is no expression change in case of before and after foraging groups (Figure 1 and 2).

**Figure 1.**
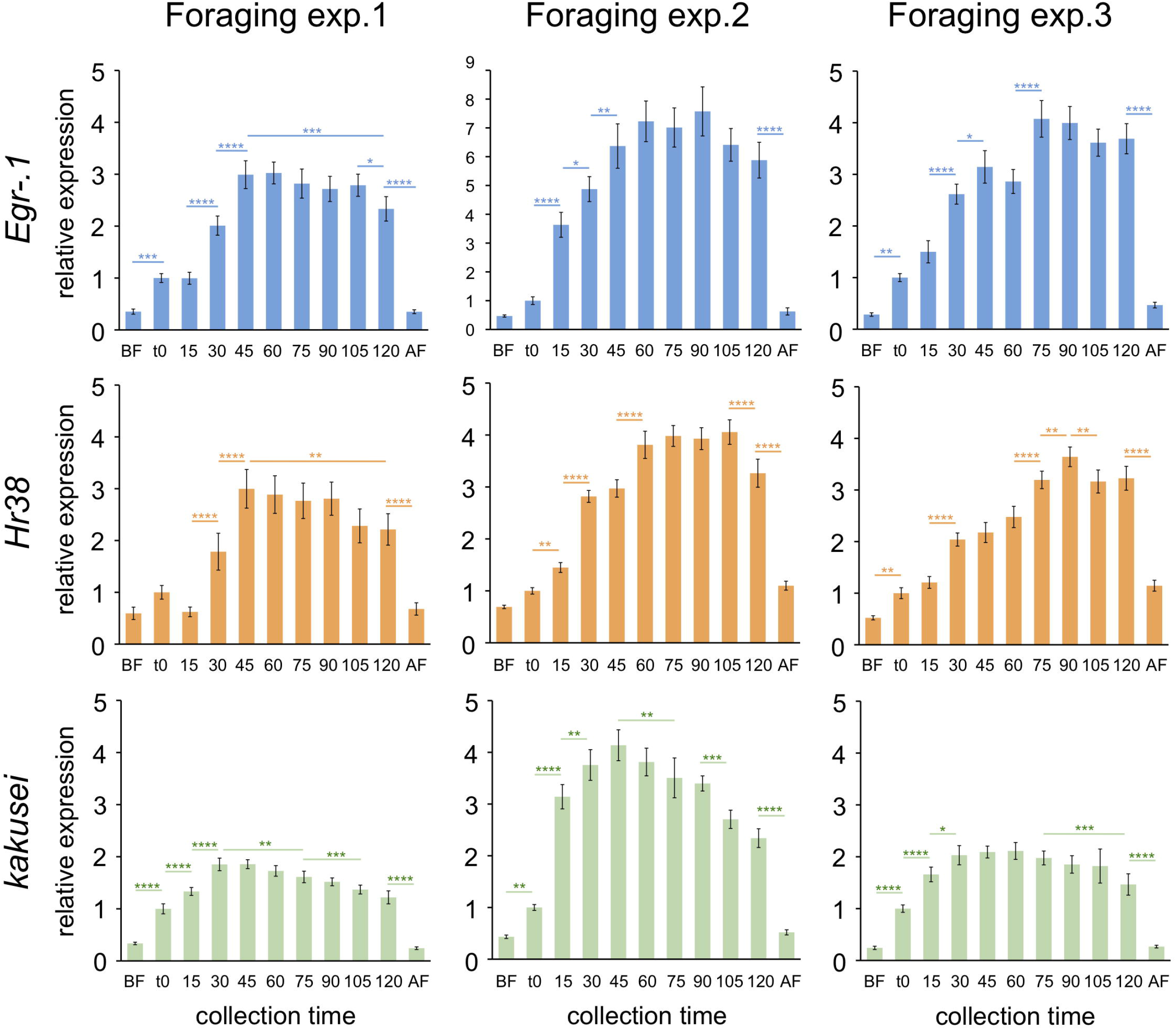
Bar graphs for three IE gene expression during, before and after foraging: The blue, orange and green represent *Egr-1, Hr38* and *Kakusei* expression during foraging at a known feeder. Data are shown as fold changes with respect to t0 (mean value was set as 1) which indicates the presentation of the feeder and start of continuous foraging. BF = before foraging and AF = after foraging. Foraging experiment 1, 2, 3 represents three replicate experiments. One-way ANOVA with Tukey-Kramer post-hoc multiple comparison: *P < 0.05, **P < 0.01, ***P < 0.001, ****P < 0.0001. Sample size for each time point is n=5. P

**Figure 2.**
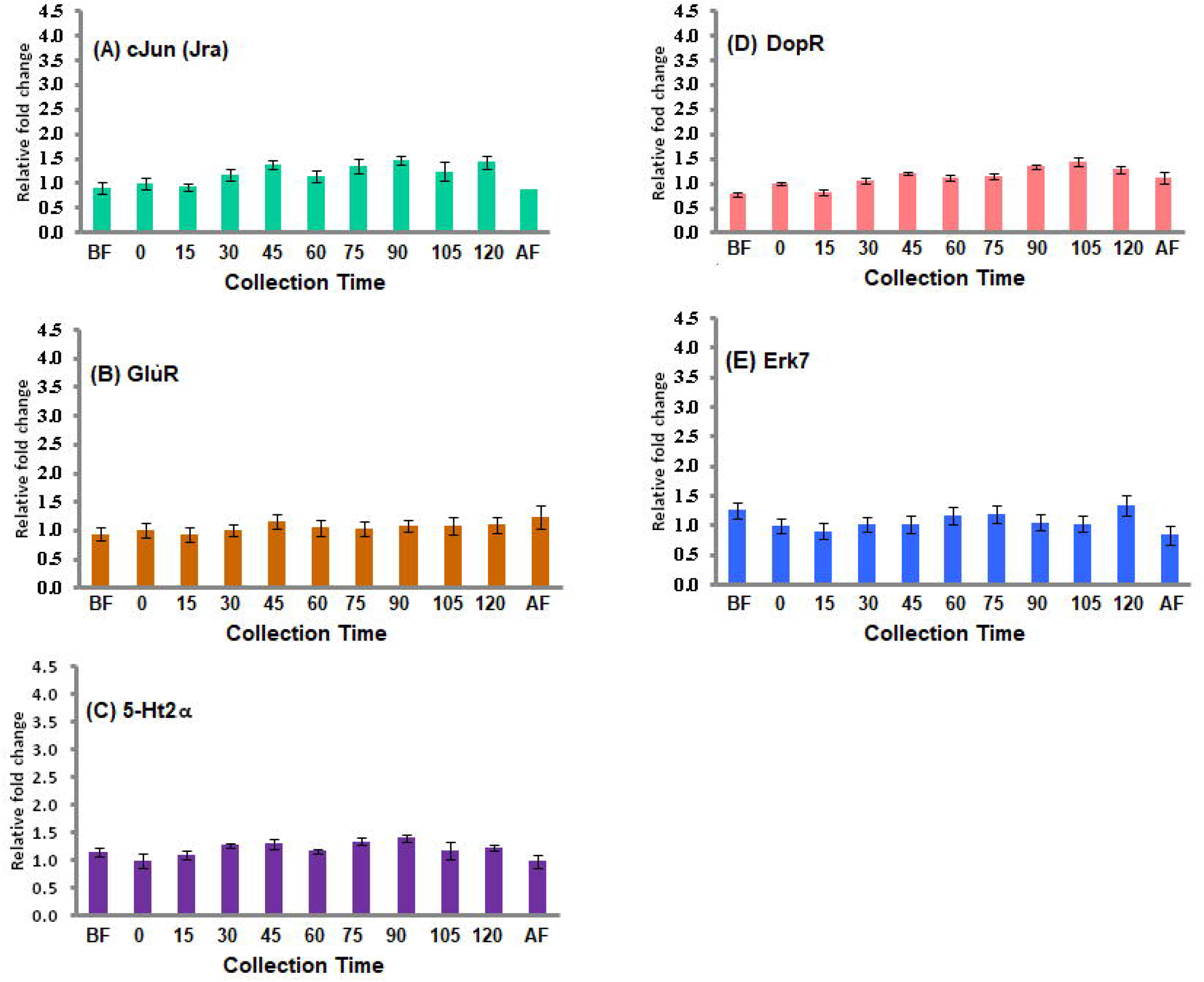
Gene expression profile for the five insignificant genes: (A) with green bars, (B) with brown bars, (C) with violet bars, (D) with red bars and (E) with blue bars represent c-Jun GluR, 5-HT2α, DopR and Erk7 gene expression during the food reward foraging. The fold changes differences were measured with respect to t0 (mean value was set as 1 at this time point). Each time point has sample size of n=5.

### *Egr-1, Hr-38* and *Kakusei* expression during foraging at the extended hour

In another experiment we collected the samples for 1 hour with 15 min intervals which is after 2 hours of reward foraging, from 16:00 hour to 17:00 hours, during which bees were continued feeding. This data have not been published before. It may be noted that the bees had been being fed since 14:00 hours, as everyday routine. This experiment was conducted because we further wanted to check if there might be any change in the gene expression during the later hour of foraging. We did not find any change in gene expression of the all the three genes *Egr-1, Hr-38* and *Kakusei*, as shown in Figure 3A. This shows that the transient over-expression is only within about the first 2 hours of reward foraging.

**Figure 3.**
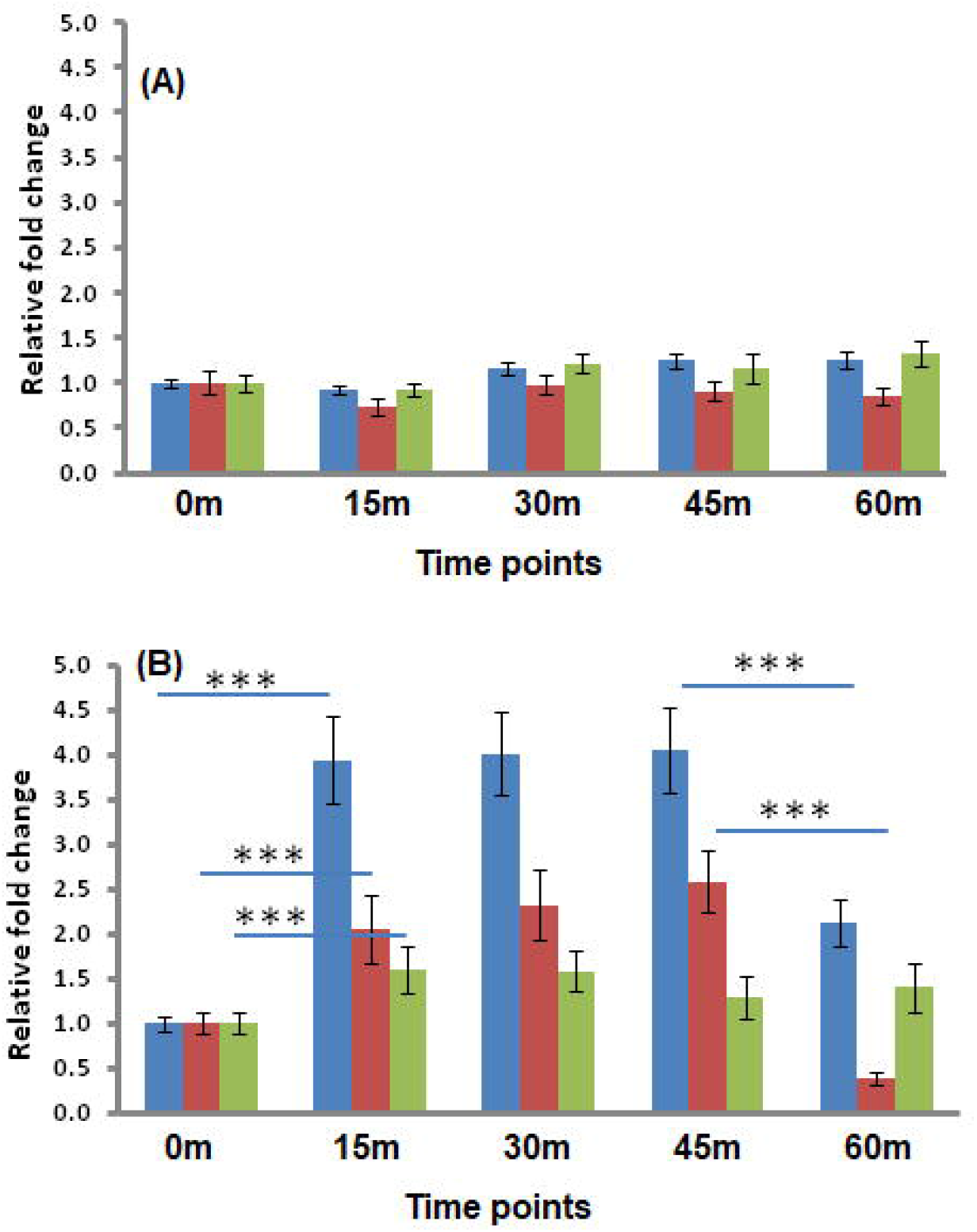
The figure 3A represents the expression profile of *Egr-1* (blue), *Hr-38* (red) and *Kakusei* (green) during the extended collection hour (15:00 hrs – 16:00 hrs). Figure 3B represents the expression of *Egr-1* (blue), *Hr-38* (red) and *Kakusei* (green) during unrewarded foraging. The fold change differences were calculateed with respect to t0 (mean value was set as 1 at this time point). Each time point has sample size of n = 5. For statistics one-way ANOVA with Turkey-Kramer post-hog multiple comparison test were performed. The number of asterisk symbol represents * P < 0.05, ** P < 0.01, *** P < 0.001, and **** P < 0.0001.

### IE genes, *Egr-1, Hr-38* and *Kakusei* expression level during unrewarded foraging

In case of unrewarded foraging the three IE genes *Egr-1, Hr38* and *Kakusei* were increased in the first 15 min of foraging upon the presentation of empty feeder plate and no further increase further, as the food was not rewarded to the bees. And gene expression level began to reduce at about 45 minutes. The summary of the result is figuratively presented in Figure 3B.

## Discussion

Our recent publications have supported that IE genes can be used as tools in searching for cellular and molecular mechanisms underlying the foraging behavior of honeybees (Singh er al 2018; Singh 2019; Singh et 2020). In those studies we investigated role of IE genes involvement during the daily of foraging and with the examination of reasonably longer duration, which was for 2 hours, during the foraging. However, we have not published in detail about method we used, basically about sample collection at different intervals in 2 hours during the daily rewarded foraging of bees. Here report the detail method with the addition of a small amount of unpublished data, shown in Figure 3A.

All these data clearly able to show that the three IE genes, *Egr-1, Hr-38* and *Kakusei* were transiently expressed during the first two hours of rewarded foraging and after which the three genes have no significant role, as indicated by their expression levels that continued to decline, even though the bees continued foraging with food reward. From these evidences, one may have an idea of choosing an appropriate time during foraging to tests other IE genes as well as their upstream or downstream molecular players and designed further experiments for finding roles they play in the specific behaviors features of honey bees during foraging. Our findings also have indicated possible role of those three IE genes in learning and memory processing and associative learning. Further research is needed in order to find the specific molecular pathways underlying those behaviors.

In other reports by Beckmann and Wilce (1997) mentioned that IE gene encoded proteins can be individually regulated in different regions of the brain depending on the type of the stimuli. This suggests that the same/different IE gene expression at different parts of the brain induced by different stimuli may signal to perform different behavioral tasks; in other words, different behaviors correspond to IE gene expression at different parts of the brain depending on the type of the stimulus. The IE genes also rapidly and transiently induced within minutes of stimulation in the absence of de-novo protein synthesis, and regulation of IEG production is necessary for the cells, because in turn it can activate the downstream target molecules that typically function as a part of a network of constitutively expressed proteins (Perez-Cadahia et al., 2011). Furthermore, different IEG reaches their pick levels at different times even though they expressed immediately after the stimulation (Bottai et al., 2002; Vazdarjanova et al., 2002). Our data agrees with the report, as in the case of three genes we investigated, *Kakusei* reached its peak earlier than *Egr-1* and *Hr-38*. These several lines of evidences clearly reveal that, it is good choice to start with IE genes when we need to start from the scratch for unwinding the molecular and cellular mechanisms underlying specific behaviors of honey bees during foraging, such as learning, memory, social interaction and social communication etc.

## Acknowledgement

Dr. Axel Brockmann is gratefully acknowledged for providing the facilities to perform and complete the experiments in his lab. National Centre for Biological Sciences (NCBS), TIFR, India, is gratefully acknowledged for the fellowships provided to Dr. Asem Surindro Singh in successfully completing this project.

## Authors’ contribution

Dr. Asem Surindro Singh and Mrs. Machathoibi Chanu Takhellambam equally contributed in this manuscript.

## Ethical statement

This work is not related to any ethical issues.

